# P4HA2-induced prolyl hydroxylation suppresses YAP1-mediated prostate cancer cell migration, invasion, and metastasis

**DOI:** 10.1101/2021.01.24.428024

**Authors:** Ming Zhu, Ruiqing Peng, Xin Liang, Zhengdao Lan, Ming Tang, Pingping Hou, Jian H. Song, Celia Sze Ling Mak, Jiwon Park, Shui-er Zheng, Ailing Huang, Xingdi Ma, Ruidong Chen, Qing Chang, Christopher J. Logothetis, Abhinav K. Jain, Sue-Hwa Lin, Hiroyuki Katayama, Samir M. Hanash, Guocan Wang

**Author notes:** These authors contributed equally to the work (Ming Zhu, Ruiqing Peng). Correspondence: Guocan Wang.

## Abstract

Yes-associated protein 1 (YAP1), a key player in the Hippo pathway, has been shown to play a critical role in tumor progression. However, the role of YAP1 in prostate cancer cell invasion, migration, and metastasis is not well defined. Through functional, transcriptomic, epigenomic, and proteomic analyses, we showed that prolyl hydroxylation of YAP1 plays a critical role in the suppression of cell migration, invasion, and metastasis in prostate cancer. Knockdown (KD) or knockout (KO) of *YAP1* led to an increase in cell migration, invasion, and metastasis in prostate cancer cells. Microarray analysis showed that the EMT pathway was activated in *Yap1*-KD cells. ChIP-seq analysis showed that YAP1 target genes are enriched in pathways regulating cell migration. Mass spectrometry analysis identified P4H prolyl hydroxylase in the YAP1 complex and YAP1 was hydroxylated at multiple proline residues. Proline-to-alanine mutations of YAP1 isoform 3 identified proline 174 as a critical residue, and its hydroxylation suppressed cell migration, invasion, and metastasis. KO of *P4ha2* led to an increase in cell migration and invasion, which was reversed upon *Yap1* KD. Our study identified a novel regulatory mechanism of YAP1 by which P4HA2-dependent prolyl hydroxylation of YAP1 determines its transcriptional activities and its function in prostate cancer metastasis.

## INTRODUCTION

Yes-associated protein 1 (YAP1), a key transcriptional coactivator in the Hippo pathway, is an important driver in cancer development and progression (1). Although YAP1 plays an oncogenic role in various cancer types, multiple studies also support a tumor-suppressive function for YAP1 in head and neck (2), breast (3–5), hematological (6), and colorectal (7, 8) cancers. Thus, the functions of YAP1 are likely context-dependent (9). YAP1 was shown to be overexpressed in prostate adenocarcinoma (PCa) and associated with cell proliferation and invasiveness in castration-resistant prostate cancer (CRPC) (10–12). However, YAP1 was found to be downregulated in the highly aggressive NEPC subset (13). Importantly, *YAP1* deletion (heterozygous and homozygous) and mutation were observed in ~3.6% of prostate cancers and was strongly associated with metastasis (**Suppl. Fig. 1A-B & Suppl. Table 1-3**). On the contrary, deletion/mutation of TAZ, another transcriptional coactivator in the Hippo pathway, was not significantly different between the primary and metastatic PCa (**Suppl. Fig. 1B**). On the cellular level, YAP1 promotes prostate cancer cell proliferation through cell-autonomous and non-autonomous mechanisms (11, 12, 14, 15), but its role in prostate cancer metastasis is not clearly defined.

Post-translational modification of YAP1, such as phosphorylation, has also been shown to regulate YAP1 cellular localization, stability, and activities (16, 17). Interestingly, we found that YAP1 proteins in prostate cancer cells are modified by proline hydroxylation, an important post-translational modification that modulates protein folding and stability in mammalian cells (18, 19). Proline hydroxylation is induced by prolyl hydroxylases, such as prolyl hydroxylase domain proteins (PHD) and collagen prolyl 4-hydroxylase (P4H). Whether YAP1 is subjected to proline hydroxylation was previously unknown, as were the effects of such modification on YAP1 function. In this study, we identified a surprising role for YAP1 in the suppression of cell migration, invasion, and metastasis in prostate, pancreatic, and breast cancers. We found that YAP1 interacts with the P4H complex and is hydroxylated at multiple proline residues. The status of proline hydroxylation of YAP1 determines its oncogenic activity in regulating cell migration, invasion, and metastasis in prostate cancer and possibly in other cancer cell types.

## RESULTS

### YAP1 suppresses cancer cell migration, invasion, and metastasis

Previously, we showed that YAP1 was highly expressed in primary tumors from the metastatic Pten/Smad4 prostate conditional knockout (KO) model (12). We first examined the effect of *Yap1* knockdown (KD) on cell migration, invasion, and metastasis in a highly metastatic Pten/Smad4-deficient CRPC cell line (referred to as PS cells hereafter) (20). Surprisingly, *Yap1* KD and *Yap1* KO led to a significant increase in cell migration and invasion (**Fig. 1A–C & Suppl. Fig. 2A**). Of note, *Yap1* KD or KO did not have a significant effect on cell proliferation as measured by the total number of cells at the end of the assays (**data not shown**). Furthermore, re-expression of human *YAP1* in *Yap1*-KO cells suppressed cell migration and invasion (**Fig. 1D & Suppl. Fig. 2B**). *YAP1*-KD in C4-2b, IGR-CaP1, and PC3 cells similarly increased migration and invasion (**Fig. 1E–G & Suppl. Fig. 2C**). However, *YAP1* KD in DU145 led to a decrease in cell migration (**Suppl. Fig. 2D**). Also, we examined whether YAP1 also suppressed cell migration in other metastatic cancers in which YAP1 has been implicated to play an important role in tumor progression (21–25). We found that *Yap1* KO or KD led to increased cell migration in iKPC mouse pancreatic cancer cells (26) and MDA-MB-231 human breast cancer cells (**Fig. 1H–I**) but not in SYO-1 synovial sarcoma cells (**Suppl. Fig. 2E**). Moreover, *Yap1* KD in PS cells promoted lung metastasis (**Fig. 1J**). Taken together, our data suggest that YAP1 suppresses cell migration, invasion, and metastasis in multiple cancer types.

**Figure 1.**
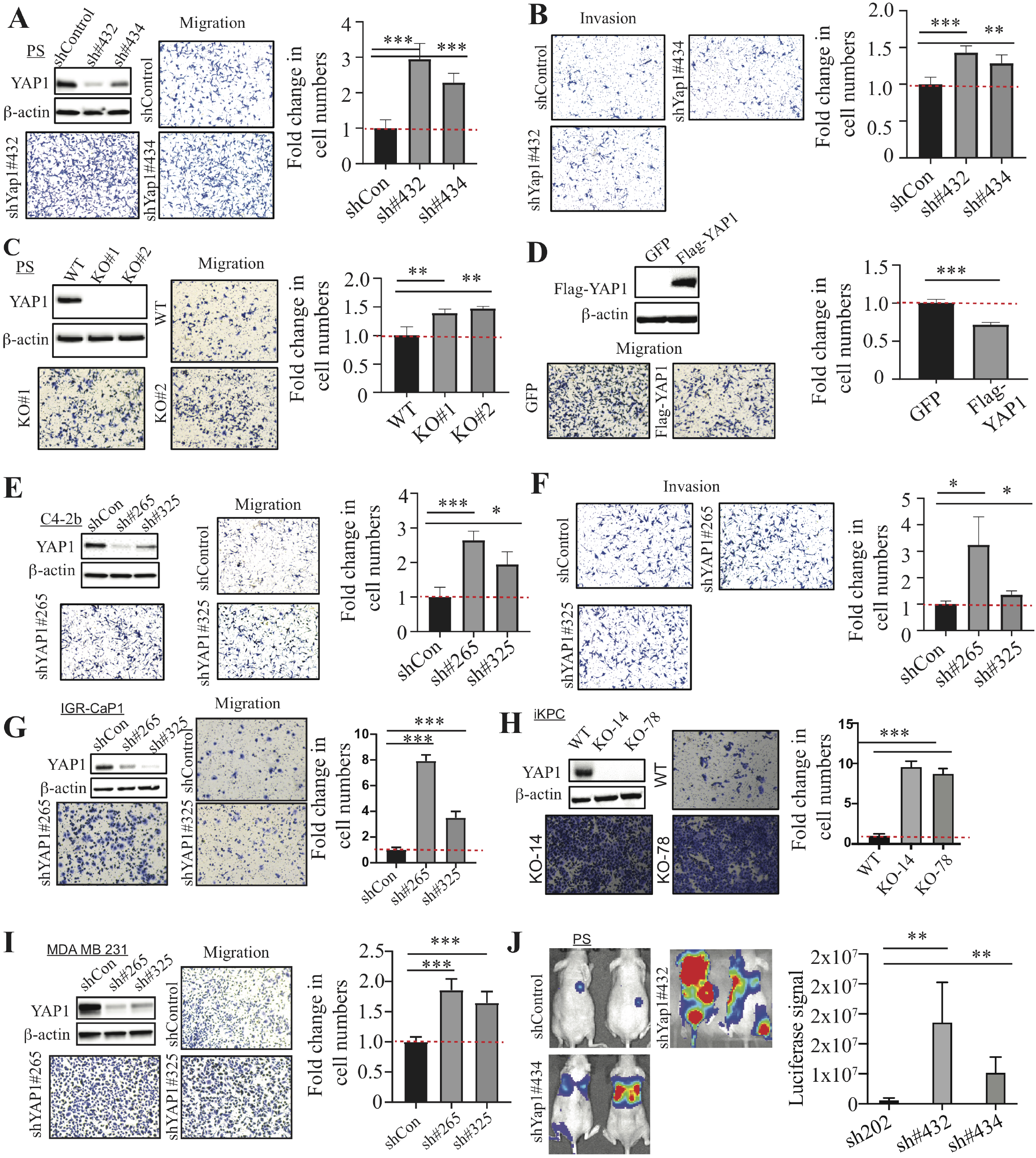
YAP1 suppresses cell migration, invasion, and metastasis. (**A-B**) Cell migration and invasion assay using PS cells transduced with control shRNA and *Yap1* shRNAs. The *Yap1* KD efficiency was confirmed by WB analysis. (**C**) Cell migration assay using *Yap1*-WT PS cells and *Yap1*-KO cells. WB analysis confirmed the KO of YAP1 expression. (**D**) Cell migration assay using *Yap1*-KO cells with GFP overexpression and YAP1 overexpression. WB analysis confirmed the overexpression of YAP1 in *Yap1*-KO PS cells. (**E-F**) Cell migration and invasion assay using C4-2b cells transduced with control shRNA and *YAP1* shRNAs. The *YAP1* KD efficiency was confirmed by WB analysis. (**G-I**) Cell migration assay using IGR-CaP1 (G), iKPC (H), and MDA-MB-231 (I) with *YAP1* KD or KO compared to control cells. (**J**) Luciferase imaging in mice injected with PS cells transduced with control shRNA and *Yap1* shRNAs through the tail vein.

### YAP1 interacts with the prolyl 4-hydroxylase complex, and its proline residues are hydroxylated

To understand the mechanisms by which YAP1 suppresses cell migration, invasion, and metastasis, we performed microarray analysis of RNA isolated from *Yap1*-KD and control PS cells (**Suppl. Table 4**). As expected, gene set enrichment analysis (GSEA) (27) identified epithelial to mesenchymal transition (EMT) as the top pathway activated in *Yap1*-KD cells (**Fig. 2A**). We also confirmed that the expression of several EMT genes, including *Postn*, *Cdh11*, *Acta2*, *Rgs4*, and *Mgp*, were upregulated upon *Yap1* KD (**Suppl. Fig. 3A**).

**Figure 2.**
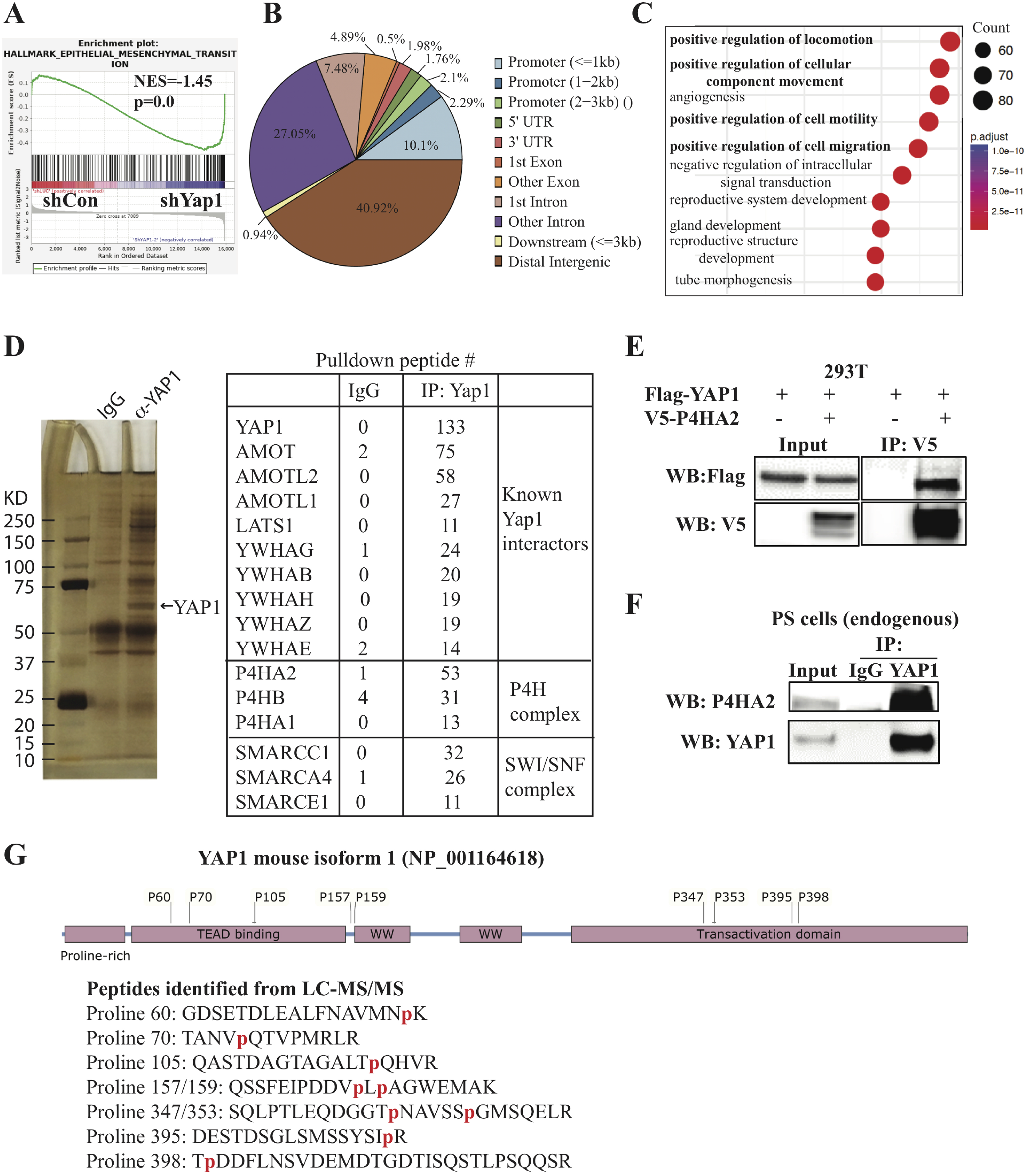
Microarray, ChIP-seq, and immunoprecipitation-mass spectrometry analyses. (**A**) GSEA analysis of microarray data from PS cells transduced with doxycycline-inducible *Yap1* shRNA identified EMT as the top pathway activated in *Yap1*-KD cells. (**B-C**) ChIP-seq analysis identified YAP1 binding sites and YAP1-regulated pathways. (**D**) Immunoprecipitation-mass spectrometry analysis identified known YAP1-interacting proteins and novel YAP1-interacting proteins. (**E**) Exogenous YAP1 interacts with exogenous P4HA2 when overexpressed in 293T cells by transfection of the indicated plasmids for co-immunoprecipitation experiments. (**F**) Endogenous YAP1 interacts with endogenous P4HA2 in PS cells. (**F**) Multiple prolyl hydroxylation sites were identified in peptides of YAP1 isoform 3 from the LC-MS/MS analysis.

Since YAP1 acts as a transcriptional coactivator, we sought to determine whether these upregulated EMT genes are direct target genes of YAP1 using ChIP-seq in PS cells. We found that YAP1 binds mostly to the distant intergenic region and other introns (**Fig. 2B**), which is consistent with previous reports (28). Pathway analyses of the top 2000 YAP1 binding sites using Gene Ontology (GO) and the Kyoto Encyclopedia of Genes and Genomes (KEGG) identified multiple pathways related to cell migration as top pathways and the expected “Hippo signaling pathway” (**Fig. 2C & Suppl. Fig. 3B**). Motif analysis using Homer (29) identified TEAD1 and AP1 motifs (**Suppl. Fig. 3C & data not shown**), which is consistent with the known physical and functional interaction between YAP1, TEAD, and AP1 (28). Importantly, we found that 194 upregulated genes and 247 downregulated genes in *Yap1* KD cells were among the top 6000 YAP1-target genes predicted by Cistrome-GO (30) (**Suppl. Table 5-7**). Among these genes, YAP1 binds to the distant intergenic region upstream or downstream of several genes upregulated in *Yap1*-KD cells (e.g., *Postn*, *Cdh11*, *Acta2*, *Rgs4*, and *Mgp*) (**Suppl. Fig. 3D & data not shown**), suggesting that YAP1 directly represses their expression. Additionally, we confirmed the binding of YAP1 to its known target genes, including *Cxcl5* and *Ccnd1* (**Suppl. Fig. 3E**).

Since the functions of YAP1 are regulated through its interacting partners, as well as by post-translational modifications (16, 17), we performed immunoprecipitation (IP) of YAP1 in PS cells followed by liquid chromatography-tandem mass spectrometry (LC-MS/MS) to identify YAP1-interacting proteins and novel post-translational modifications (**Fig. 2D**). As expected, we identified multiple proteins previously shown to interact with YAP1, such as AMOT and the SWI/SNF complex (**Fig. 2D**). Interestingly, proteins of P4H complex (P4HA1, P4HA2, and P4HB) were identified among the top YAP1-interacting proteins (**Fig. 2D**). On the contrary, prolyl 3-hydroxylase was not identified in the IP-MS (**data not shown**). Collagen P4H, an α2β2 tetrameric complex, specifically catalyzes 4-hydroxylation of proline (18, 19) through its catalytic α subunit (P4HA). Because P4HA2 was the most abundant protein of the P4H complex pulled down by YAP1, we focused on its interaction with YAP1. We showed that overexpressed Flag-YAP1 efficiently pulled down overexpressed P4HA2 in 293T cells (**Fig. 2E**). Also, endogenous YAP1 was found to interact with P4HA2 in PS cells (**Fig. 2F**).

Give the proline hydroxylase activity of the P4H complex (18, 19), we examined whether proline residues in YAP1 were hydroxylated. We identified nine hydroxylated proline residues (proline 60, 70, 105, 157, 159, 347, 353, 395, 398) in mouse YAP1, eight of which were evolutionarily conserved between mouse and human (**Fig. 2G & Suppl. Fig. 4A-B**), suggesting that they might play a role in regulating YAP1 functions.

### Hydroxylation at proline 174 of YAP1 plays a critical role in suppressing cell migration, invasion, and metastasis

Given the critical regulatory roles of proline hydroxylation in proteins (18, 19), we decided to examine whether proline hydroxylation modulates YAP1 functions. We first generated hydroxylation-defective human YAP1 mutants by mutating proline to alanine (Mut1-3: P75/85/120A; Mut4-5: P172/174A; Mut6-9: P348/352/394/397A) (**Fig. 3A & Suppl. Fig. 4B**). These YAP1 mutants were overexpressed in *Yap1*-KO PS cells to avoid the possible interference of the endogenous wild type (WT) YAP1. The expression of all the YAP1 mutants was similar but higher than the WT (**Fig. 3B**). We did not observe any significant difference in cell growth *in vitro* and *in vivo* (**Suppl. Fig. 5A-B**) between YAP1 WT and mutants. However, we found that there is an increase in cell migration and invasion in Mut4-5–overexpressing cells compared to WT-overexpressing cells (**Fig. 3C–D**), suggesting that mutations of proline 172 and 174 to alanine abolished the activity of YAP1 in suppressing cell migration and invasion. Consistent with the *in vitro* findings, PS cells with YAP1 Mut4-5 overexpression increased the colonization of cancer cells in the lung compared to WT, Mut1-3, and Mut6-9 (**Fig. 3E & Suppl. Fig. 5C**).

**Figure 3.**
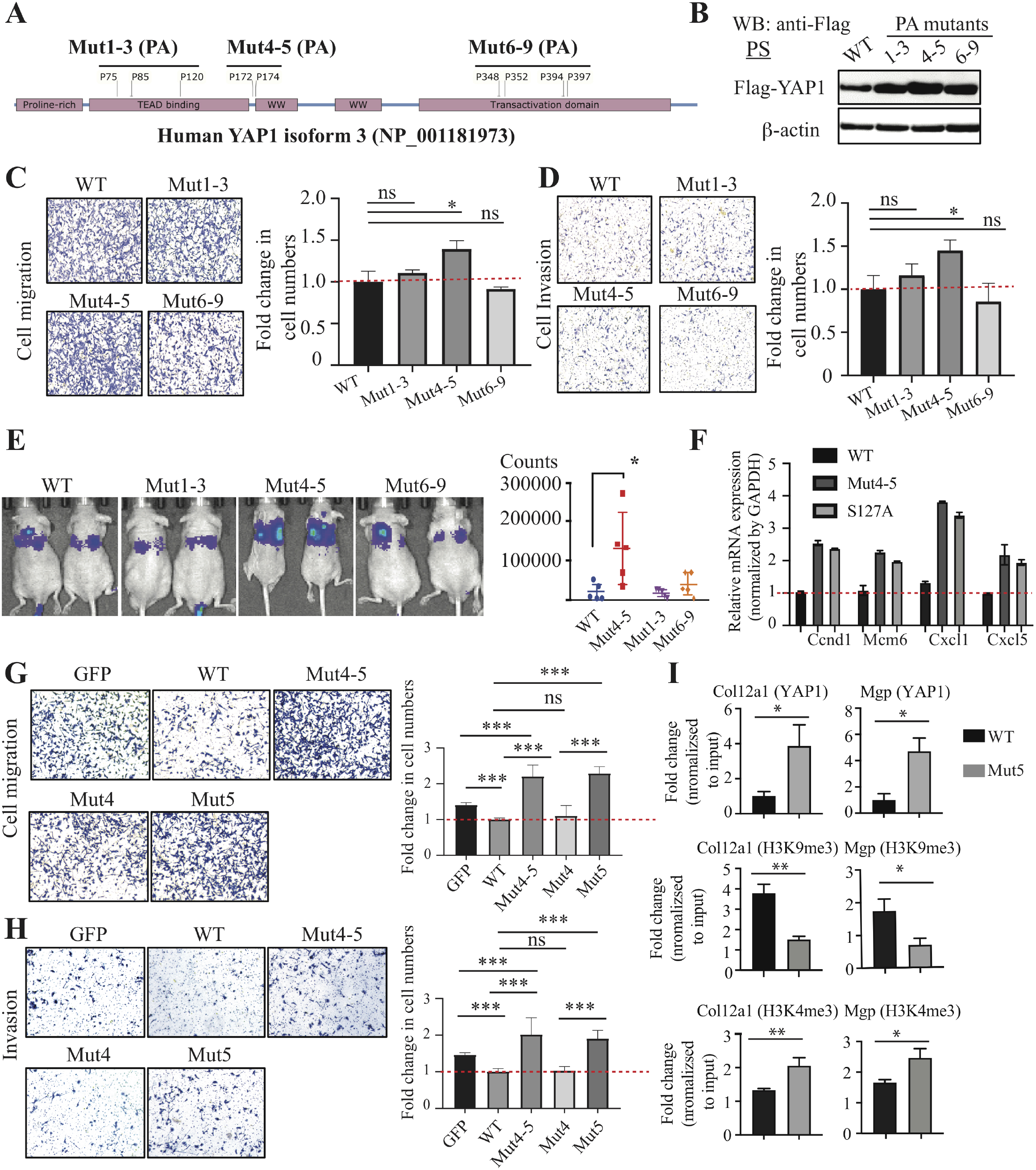
Prolyl hydroxylation of YAP1 suppressed cell migration, invasion, and metastases. (**A**) Scheme showing the strategy to generate prolyl hydroxylation–defective YAP1 mutants by mutating proline to alanine (PA): Mut1-3 (P75/85/120A), Mut4-5 (P172/174A), and Mut6-9 (P348/352/394/397). Human YAP1 isoform 3 was used. (**B**) Expression of YAP1 WT and PA mutants in *Yap1*-KO PS cells. (**C-D**) Cell migration and invasion assay in *Yap1*-KO PS cells with overexpression of YAP1 WT and PA mutants. (**E**) Tail vein injection of *Yap1*-KO cells with overexpression of YAP1 WT and PA mutants. (**F**) qPCR analysis of YAP1 target genes in *Yap1*-KO PS cells with overexpression of YAP1 WT, Mut4-5, and the constitutively active S127A mutant. (**G-H**) Cell migration and invasion assay using *Yap1*-KO PS cells with overexpression of GFP, YAP1 WT, Mut4-5, Mut4, and Mut5. (**I**) ChIP-qPCR analysis of YAP1, H3K4me3, and H3K9me4 binding sites in *Col12a1* and *Mgp.*

To determine whether prolyl hydroxylation controls the transcriptional activities of YAP1, we examined the expression of YAP1 target genes *Ccnd1*, *Mcm6*, *Cxcl1*, and *Cxcl5* in *Yap1*-KO cells that overexpressed YAP1 WT, Mut4-5, or the constitutively active S127A mutant (31). Both Mut4-5 and the S127A mutant dramatically increased the expression of these genes compared to the YAP1 WT (**Fig. 3F & Suppl. Fig. 5D**), suggesting that the non-hydroxylated YAP1 is more transcriptionally active than the hydroxylated YAP1.

To further pinpoint which proline residue of YAP1 is critical for its function in suppressing cell migration and invasion, we generated site-specific proline-to-alanine mutants of human YAP1 (P172A and P174A mutants), which corresponded to proline 157 and 159 in mouse YAP1. We overexpressed GFP control, YAP1 WT, and YAP1 mutants (Mut4-5: P172/174A; Mut4: P172A; Mut5: P174A) in *Yap1*-KO cells (**Suppl. Fig. 5E**) and examined their effects on cell migration and invasion. We found that both Mut4-5 and Mut5, but not Mut4, dramatically increased cell migration and invasion compared to the GFP control (**Fig. 3G–H)**, suggesting proline 174 is the critical hydroxylation site that regulates YAP1 activity in cell migration and invasion. Importantly, overexpression of Mut4-5 and Mut5 similarly increased the cell migration of PC3 and TRAMPC2 cells (**Suppl. Fig. 5F**). We then performed ChIP-qPCR to determine whether the increased expression of YAP1 target genes in Mut4-5 expressing cells is due to increased YAP1 binding to chromatin. We found that Mut5 binding to the promoter/distal enhancers of its target genes was significantly increased compared to WT (**Fig. 3I & Suppl. Fig. 5G**). To examine the transcriptional activation and repression of YAP1 target genes, we examined H3K9me3, a mark associated with transcriptional repression (32), and H3K4me3, a hallmark of active chromatin enriched at active promoters and correlates with transcriptional activity (33), in the promoters/enhancers of YAP1 target genes. We found that H3K9me3 was significantly decreased and H3K4me4 was significantly increased in the regulatory region of several YAP1 target genes (e.g., *Col12a1*, *Mgp*, *Postn*, *Cxcl12*) (**Fig. 3I & Suppl. Fig. 5G**). Taken together, our data suggest that hydroxylation at proline 174 of YAP1 plays a critical role in regulating cell migration, invasion, and metastasis by repressing the expression of a subset of its target genes.

### Loss of P4HA2 promotes cell migration and invasion through YAP1

Given the observed hydroxylation of YAP1 in PS cells (**Fig. 2G**) and the effect of hydroxylation-defective YAP1 mutants on migration and invasion (**Fig. 3C–H & Suppl. Fig. 5F**), the ability of YAP1 to suppress cell migration and invasion appeared to be regulated by the P4H complex. We examined the effect of *P4ha2* KO on cell migration and invasion and found that *P4ha2* KO significantly increased cell migration and invasion compared to WT cells (**Fig. 4A–C**), which was not due to an increase in cell proliferation (**data not shown**). Interestingly, we found that YAP1 target genes *Postn*, *Col12a1*, *Mgp, Ccl5, and Cxcl12* were significantly upregulated in *P4ha2-*KO cells compared to control cells (**Fig. 4D & data not shown**), suggesting YAP1 is transcriptionally more active upon loss of P4HA2. Importantly, KD of *Yap1* in *P4ha2*-KO cells abolished the effect of *P4ha2* KO on cell migration and invasion (**Fig. 4E–F & Suppl. Fig. 5H**). Taken together, our data indicate that P4HA2 suppresses cell migration and invasion through proline hydroxylation of YAP1.

**Figure 4.**
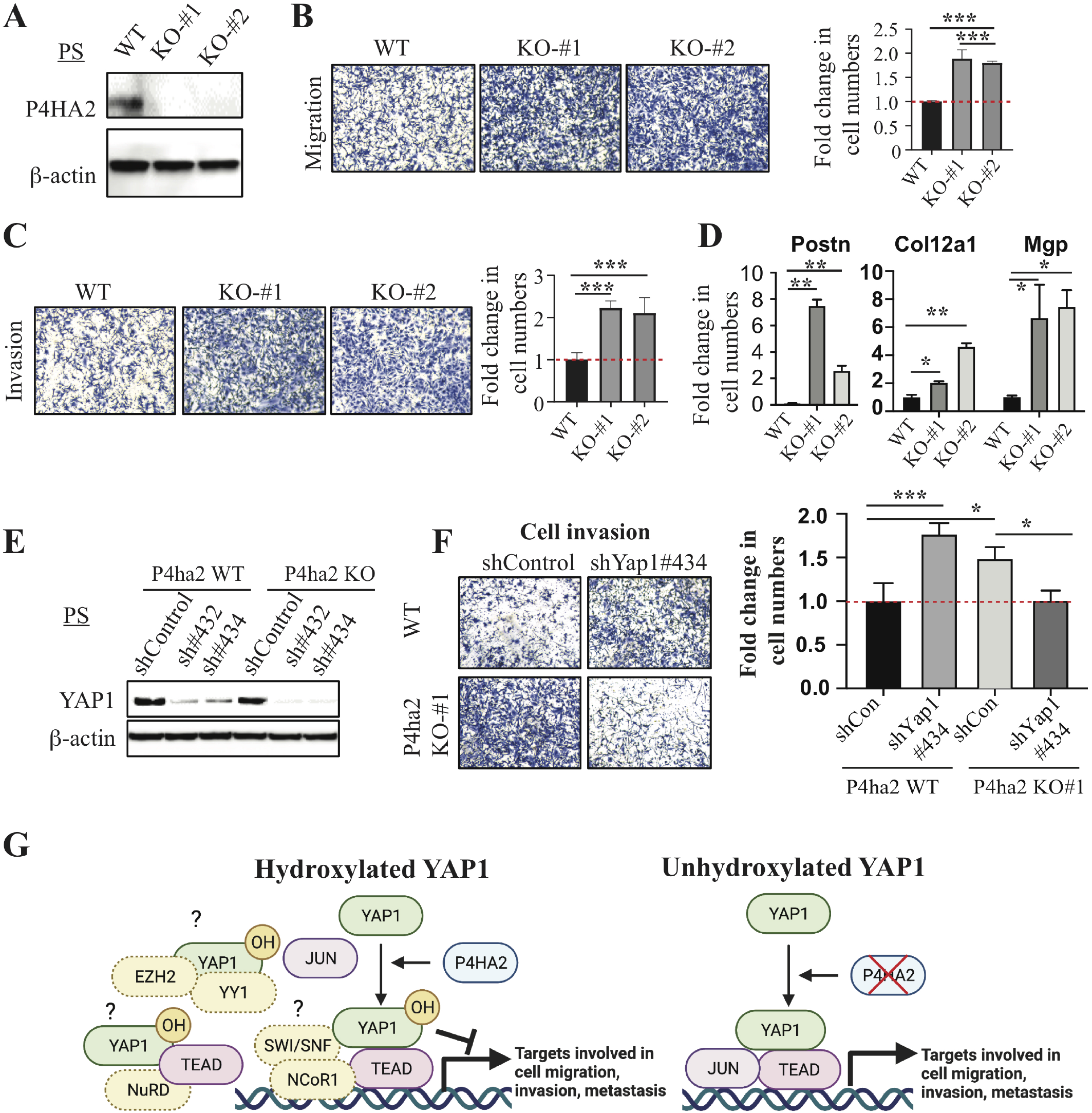
P4HA2 suppresses cell migration and invasion through Yap1. (**A**) WB analysis of P4HA2 in *P4ha2*-WT and *P4ha2*-KO PS cells. (**B-C**) Cell migration and invasion using *P4ha2*-WT and *P4ha2*-KO PS cells. (**D**) qPCR analysis of YAP1 target genes (*Postn, Col12a1, and Mgp*) in *P4ha2* KO and WT cells. (**E**) WB analysis of YAP1 in *P4ha2*-WT and *P4ha2*-KO PS cells transduced with Yap1 shRNAs. (**F**) Cell invasion assay in *P4ha2*-WT and *P4ha2*-KO PS cells transduced with shYap1#434. (**G**) A model for hydroxylation-dependent YAP1 function in cell migration, invasion, and metastasis (Created with BioRender.com). Left: P4HA2-mediated hydroxylation of YAP1 may impair its interactions with transcription factors such as JUN or enhance the recruitment of corepressor, such as SWI/SNF-NCoR1, NuRD, and EZH2/YY1, which results in a decrease in the expression of genes involved in cell migration, invasion, and metastasis. Right: In the absence of P4HA2, non-hydroxylated YAP1 may efficiently interact with transcription factors such as JUN to activate genes involved in cell migration, invasion, and metastasis.

## DISCUSSION

In contrary to previous findings that KD of *YAP1* in LNCaP-C4-2 cells impaired cell migration and invasion, we demonstrated that KD or KO of *Yap1* in mouse (PS and TRAMPC2) and human (C4-2b, IGR-CaP1, and PC3) prostate cancer cells led to enhanced cell migration, invasion, and metastasis. The discrepancy between our study and the previous one is not clear. Our data also showed that *YAP1* KD in DU145 cells suppressed cell migration whereas YAP1 KD/KO promoted cell migration in iKPC cells and MDA-MB-231 cells. These findings strongly suggest that YAP1 plays a context-dependent function in cell migration, invasion, and metastasis. Further studies are necessary to define molecular basis underlying the context-dependent functions of YAP1 in cell migration, invasion, and metastasis. Importantly, the clinical significance of YAP1 loss in PCa patients was supported by the strong association of YAP1 deletion with metastatic PCa and the loss of YAP1 protein in advanced PCa (12, 13). Loss of YAP1 function via post-translational modification by proline hydroxylation will be also an important mechanism of clinical significance. Furthermore, our unpublished data showed that TAZ similarly suppresses cell migration and regulates a common set of genes as YAP1, suggesting functional redundancy between YAP1 and TAZ.

Mechanistically, our data suggest that prolyl hydroxylation plays an important role in the regulation of YAP1 activities, which can both activate and repress transcription (**Fig. 4G**). In P4HA2 WT cells, YAP1 may suppress gene expression through its interaction with SWI/SNF complex, which was identified as YAP1-interating proteins in our study and has been shown to regulate both activation and repression of the same promoters (34, 35), in part through corepressor NCoR1 (36). Also, YAP1 may recruit YY1 and EZH2 (37) or recruit the NuRD complex to suppress the expression of its target genes (38). Interestingly, proline 174 is within the first WW domain of YAP1, which is crucial for the transcriptional activities of YAP1 through its interaction with transcription factors that contain PPxY motifs (39). Thus, our findings suggest that prolyl hydroxylation at P174 of YAP1 may impair its interaction with key transcription [e.g., c-JUN (28), TEAD], resulting in a decrease in both binding to its target gene and reduced transcriptional activation. This notion is supported by our findings that YAP1-Mut5 binds more efficiently to its target genes. YAP1 hydroxylation may also be required for the efficient recruitment of the transcription corepressor complexes (e.g., NCoR1, NuRD), as our data showed that YAP1 P174A OE increased H3K4me3 and reduced H3K9me3 for a subset of YAP1 target genes. Thus, our data suggest that P174A YAP1 mutant not only has increased binding to its target genes, but also induces a switch from repressive transcription to active transcription for a subset of genes. Further studies are needed to delineate the effect of prolyl hydroxylation of YAP1 on the dynamics of the epigenomic landscape. YAP1 is known to be regulated by phosphorylation at multiple serine residues (40). Our studies shed light on a new aspect of YAP1 regulation, which may also be involved in many YAP1-mediated cellular activities yet to be identified.

The P4H α subunit (P4HA) has three isoforms (P4HA1-3) in mammalian cells (19). Since P4HA1 and P4HA2 were both identified as YAP1-interacting proteins in our MS analysis, P4HA1 may also suppress cell migration, invasion, and metastasis through prolyl hydroxylation of YAP1. However, P4HA1 was previously shown to promote prostate cancer progression (41), and further studies are needed to clarify the role of P4HA1 in regulating YAP1 functions and cell migration, invasion, and metastasis. Since LS-MS mass spectrometry analysis cannot distinguish 3-prolyl hydroxylation from 4-prolyl hydroxylation, we cannot rule out the presence of 3-hydroxyl proline in YAP1. Given that P3H1 was not identified as YAP1-interacting proteins, the high specificity of the P4H and P3H towards prolyl hydroxylation strongly indicates that the hydroxyproline identified in YAP1 is 4-hydroxyproline. Future experiments combining liquid chromatography retention time differences with mass spectrometry using ETD-HCD fragmentation, complemented by ab initio calculations are needed to address this issue (42).

Although collagen deposition is generally associated with tumor progression and invasive behavior (43), it also plays a tumor-suppressive role (44). P4HA1 is the major isoenzyme in most cells, and *P4ha1*−/− leads to embryonic lethality in mice due to abnormal deposition of collagen IV (45). In contrast, *P4ha2*−/− mice had no apparent abnormalities (46). Given that we did not observe any significant difference in the expression of P4HA1 between *P4ha2* WT and KO cells (**Suppl. Fig. 5I**), collagen deposition in *P4ha2* KO/KD cells may not be significantly impacted. Given the role of P4HA2 in promoting secretion and deposition of collagen and cell invasion in breast cancer (47, 48), the seemly contradictory findings on the differential effect of *P4ha2* KO/KD and collagen deposition on cell migration may warrant further studies. Furthermore, due to the lack of an antibody that can specifically recognize the hydroxylated proline 174 of YAP1, we cannot assess the clinical relevance of prolyl hydroxylation of YAP1 in tumor samples from prostate cancer patients. The development of such antibodies is warranted and would allow us to examine the association of prolyl hydroxylation of YAP1 and P4HA2 expression in prostate cancer specimens.

In summary, our findings support a model in which P4HA2-mediated prolyl hydroxylation serves as a molecular switch that controls the activities of YAP1 in cell migration, invasion, and metastasis (**Fig. 4G**).

## MATERIALS AND METHODS

The reagents and assays as well as bioinformatic/statistical analyses were described in Supplementary Information.

## ACKNOWLEDGEMENTS

We thank Ronald DePinho and Haoqiang Ying for helpful discussions, Sarah E. Townsend for editing, and Dr. Timothy C. Thompson for providing RM-1 cells. G.W. is supported by the funding from MDACC (Moon Shot, IRG, and PCRP), UT STARs Award, NIH R00 CA194289 and P50 CA140388 as well as NIH P30CA016672 for the use of Research Animal Support Facility, Flow Cytometry and Cellular Imaging Core Facility, and Functional Genomics Core.

## CONTRIBUTIONS

MZ and GW contributed to the study’s conception and design of this study. GW, MZ, RP, XL, ZL, MT, PH, JHS, CSM, JP, SZ, AH, XM, RC, QC, AKJ, and HK performed the experiments and acquired, analysed, and interpreted the data (e.g., statistical analysis, biostatistics, computational analysis). CJL, SMH, and GW contributed to the supervision of the project. SHL and GW contributed to the writing and editing the manuscript.

