## Supplementary material for "P4HA2-induced prolyl hydroxylation suppresses YAP1-mediated prostate cancer cell migration, invasion, and metastasis": Suppl. Materials and Methods

**SUPPLEMENTARY MATERIALS AND METHODS**

**Western Blotting**

Cells were lysed on ice using RIPA buffer (Boston BioProducts) supplemented with Protease and Phosphatase Inhibitor Cocktail (Thermo Fisher Scientific). Proteins with loading buffer (20 μg) were subjected to SDS-PAGE and transferred onto a nitrocellulose membrane. Then, the membrane was incubated with primary antibodies for 16 h at 4°C and washed 3 times with Tris- buffered saline (TBS) containing 0.1% Tween 20. Next, the membrane was incubated with HRP- conjugated secondary antibody for 2 h at room temperature. After washing 3 times with TBS containing 0.1% Tween 20, the membrane was exposed to an ECL solution and imaged with an Azure Biosystems c600. The following antibodies were used in this study: Anti-Yap antibody (Cell Signaling Technology, #14074), Anti-Beta Actin antibody (Thermo Fisher Scientific, MA5- 15739), Anti-Flag Tag antibody (Sigma-Aldrich, F1804), Anti-V5 Tag antibody (Thermo Fisher Scientific, R960-25), Anti-P4HA2 antibody (Proteintech, 13759-1-AP), Anti-Rabbit IgG HRP- linked antibody (Cell Signaling Technology, 7074), Anti-Mouse IgG HRP-linked antibody (Cell Signaling Technology, 7076), Rabbit IgG (Cell Signaling Technology, 3900), Mouse IgG (Cell Signaling Technology, 5415), TrueBlot Anti-Rabbit IgG HRP-linked antibody (Rockland, 18- 8816-33), and TrueBlot Anti-Mouse IgG HRP-linked antibody (Rockland, 18-8817-33).

**Quantitative RT-PCR**

Cell mRNA was isolated by RNeasy Kit (Qiagen) and reverse transcribed using Superscript III cDNA Synthesis Kit (Life Technology). Quantitative PCR was performed using SYBR-GreenER Kit (Life Technology). The primers used were as follows: mouse Ccnd1 (forward: GCGTACCCTGACACCAATCTC; reverse: ACTTGAAGTAAGATACGGAGGGC), mouse Mcm6 (forward: GCTGTTCCTAGACTTCCTGGA; reverse: CAACCAGCGTGTTTCTCTCAG), mouse Cxcl1 (forward: TGAGCTGCGCTGTCAGTGCCT; reverse: AGAAGCCAGCGTTCACCAGA), mouse Cxcl5 (forward: TCCAGCTCGCCATTCATGC; reverse: TTGCGGCTATGACTGAGGAAG), mouse Col12al (forward: AAGTTGACCCACCTTCCGAC; reverse: GGTCCACTGTTATTCTGTAACCC), mouse Postn (forward: TGGTATCAAGGTGCTATCTGCG; reverse: AATGCCCAGCGTGCCATAA), mouse Cdh11 (forward: CTGGGTCTGGAACCAATTCTTT; reverse: GCCTGAGCCATCAGTGTGTA), mouse Acta2 (forward: GTCCCAGACATCAGGGAGTAA; reverse: TCGGATACTTCAGCGTCAGGA), mouse	Rgs4 (forward: GAGTGCAAAGGACATGAAACATC; reverse: TTTTCCAACGATTCAGCCCAT), mouse Mgp (forward: GGCAACCCTGTGCTACGAAT; reverse: CCTGGACTCTCTTTTGGGCTTTA), mouse GAPDH (forward: AGGTCGGTGTGAACGGATTTG; reverse: TGTAGACCATGTAGTTGAGGTCA).

**Chromatin-immunoprecipitation and quantitative PCR (ChIP-qPCR)**

PCR primers were designed using YAP1 ChIP-seq data and ENCODE ChIP-seq data available at UCSC Genome Browser based on the YAP1, H3K4me3, and H3K9me3 peaks for the selected genes. The primers used were as following: Postn_Yap1_F: ACATGGCCCCAGTTTCATAG, Postn_Yap1_R: CTTCTGTCTCTGCCACCACA; Col12a1_Yap1_F: CCCTTGTGGTTTGCTTGTTT, Col12a1_Yap1_R:	CACCAGGGAGTGGCACTATT; Mgp_Yap1_F: GGAGAGGCTCCTACATGCTG, Mgp_Yap1_R:	CTGCCCACGCTGTGTAGATA; Cxcl12_Yap1_F:	CCTGTCTGCTGAAAGGAAGG, Cxcl12_Yap1_R: AACGCACAGCTAGAACAGCA; Postn_H3K4me3_F: TGTGTGGAGAGCGACTTTTG, Postn_H3K4me3_R: AGGCCAGGTCAGAAGAGTGA; Col12a1_H3K4me3_F: TAAAATTTTGGCGCAGCTCT, Col12a1_H3K4me3_R: CAACTGCGGACTGCATTCTA; Mgp_H3K4me3_F	: TTGGCCCAAGTACTCATTCC Mgp_H3K4me3_R: ATTCGAAAGGCAAACCTGTG; Cxcl12_H3K4me3_F: TGGGCTGCTGTTCCTACTCT, Cxcl12_H3K4me3_R: GCTGCTGAAACAGTTGTCCA; Postn_H3K9me3_F: TCATGGTACTGGCATCTCCA, Postn_H3K9me3_R: CACAAAGAGGCTGTGTTCCA; Col12a1_H3K9me3_F: CCCTTGTGGTTTGCTTGTTT, Col12a1_H3K9me3_R: CACCAGGGAGTGGCACTATT; Mgp_H3K9me3_F: ACGTTGTGCAGTTGTCCAAG, Mgp_H3K9me3_R: CATCTTCAGCCCTGCCTTAC; Cxcl12_H3K9me3_F: AAAGCCTCTAGCCCTCCAAG, Cxcl12_H3K9me3_R: TTCCTGCCTTTCCAGAAGAA.

**Liquid chromatography-tandem mass spectrometry (LC-MS/MS)**

Proteins were separated using sodium dodecyl sulfate polyacrylamide gel electrophoresis and silver stained using SilverQuest Silver Staining Kit (Cat. LC6070, Invitrogen). The respective gel lanes were cut, destained, alkylated for 30 minutes with 0.2% DTT in 100 mM NH4HCO3, additionally alkylated for 30 mins with 0.3% acrylamide in 100 mM NH4HCO3, followed by washing for 30 mins with 5% acetic acid in 50% methanol. Following aspiration of wash buffer, the gel pieces were additionally washed in 100 mM NH4HCO3 and then dried for 45 mins using a SpeedVac vacuum concentrator and subjected to overnight tryptic digestion. Digested peptides were extracted from the gel pieces adding equal volume of 0.1% TFA in H2O/75% acetonitrile mixing for 15 mins, followed by spin and supernatant harvest and a second extraction using 50 µL of a 0.1% TFA in H2O/75% acetonitrile for 15 mins. The pooled supernatants were vacuum concentrated to dryness and stored at -80°C for subsequent LC– MS/MS analysis.

The tryptic peptides were separated by reversed-phase chromatography using an EASYnano HPLC system (Thermo Scientific) coupled online with Orbitrap ELITE mass spectrometer (Thermo Scientific). Mass spectrometer parameters were spray voltage 2.5 kV, capillary temperature 320°C, FT resolution 60,000, FT target value 1×106, LTQ target value 3x104, 1 FT microscan with 500 ms injection time, and 1 LTQ microscan with 10 ms injection time. Mass spectra were acquired in a data-dependent mode with the m/z range of 300-1800. The full mass spectrum (MS scan) was acquired by the FT, and tandem mass spectrum (MS/MS scan) was acquired by the LTQ with a 35% normalized collision energy. Acquisition of each full mass spectrum was followed by the acquisition of MS/MS spectra for the five most intense +2 or +3 ions within a one-second duty cycle. The minimum signal threshold (counts) for a precursor occurring during an MS scan was set at 1000 for triggering an MS/MS scan.

The acquired LC-MS/MS data was processed by the Proteome Discoverer 1.4 (Thermo Scientific). The Mascot was used as a search engine with the parameters including cysteine (Cys) alkylated with acrylamide (71.03714@C) as a fixed modification and proline hydroxylation, methionine	(Met)	oxidation, 13C6	15N2	lysine (80.14199@Heavy labeled-K) and 13C6 15N4 arginine (10.008269@Heavy labeled-R) as variable modifications. Data was searched against the UniProt human database 2017 and further filtered with false discovery rate (FDR) ≤ 1%.

**SUPPLEMENTARY FIGURES**

**Suppl. Figure 1. YAP1 is deleted and mutated in a subset of human prostate cancers.** (**A**) YAP1 deletions and mutations in prostate cancer patient samples were obtained from cBioPortal using combined studies. (**B**) YAP1 mutation/deletion (heterozygous and homozygous) are significantly associated with prostate cancer metastasis (P< 0.0001), whereas TAZ mutation/deletion is not associated prostate cancer metastasis.

**Suppl. Figure 2. YAP1 regulates cell migration and invasion in context-dependent manner.**

(**A**) Cell invasion assay using Yap1-WT and Yap1-KO PS cells. (**B**) Cell invasion assay using Yap1-KO PS cells with overexpression of GFP or YAP1. (**C**) Cell migration and invasion assay using YAP1-KD PC3 cell and control cells. (**D**) Cell migration assay using YAP1-KD DU145 cell and control cells. (**E**) Cell migration assay using YAP1-KD SYO-1 cell and control cells.

**Suppl. Figure 3.** Microarray and ChIP-seq analysis support the role of YAP1 in the regulation of cell migration and invasion. (**A**) qPCR showing that the mRNA of YAP1 target genes is increased in Yap1-KD PS cells. (**B-C**) Homer was used to analyze ChIP-seq data to identify Yap1-regulated pathways and Yap1-binding motifs. (**D-E**) ChIP-seq peaks in selected Yap1 target genes.

**Suppl. Figure 4. LS-MS/MS analysis identified prolyl hydroxylation sites in mouse Yap1.** (**A**) The ion mass spectra of the recovered peptides of mouse Yap1 isoform 1 showed that P157 and P157 were hydroxylated. (**B**) Alignment of human and mouse YAP1 proteins showed that 8 out of the 9 hydroxylated prolines are conserved in human and mouse YAP1.

**Suppl. Figure 5. Prolyl hydroxylation of YAP1 promotes cell migration, invasion, and metastasis.** (**A-B**) YAP1 hydroxylation-defective mutants did not affect cell proliferation when they were overexpressed in Yap1-KO PS cells compared to Yap1 WT as shown by foci-forming assay (A) and subQ transplantation (B). (**C**) H & E staining of representative lung tissues from mice that received tail vein injection of Yap1-KO cells with overexpression of YAP1 WT and Mut4-5. (**D**) qPCR analysis of YAP1 target genes in PS cells with overexpression of YAP1 WT and Mut4-5. (**E**) WB analysis of YAP1 WT and Mut4-5, Mut4, and Mut5 in PS cells. (**F**) Migration assay using PC3 and TRAMPC2 cells with overexpression of YAP1 WT, Mut4-5, and Mut5. (**G**) ChIP-qPCR analysis of YAP1 target genes Postn and Cxcl12 for YAP1 binding to their promoters/enhancers and the differences in H3K4me3 and H3m9me3. (**H**) Invasion assay using *P4ha2*-KO PS cells transduced with control shRNA or shYap1#434. (I) WB analysis of P4HA1 expression in *P4ha2*-KO PS cells and *P4ha2*-WT PS cells.

**SUPPLEMENTARY TABLES**

**Supp. Table 1-3. YAP1 deletion and mutations identified in prostate cancers using cBioportal**. Table 1: combined study; Table 2: TCGA; Table 3: Abida et al. 2019 mCRPC.

**Supp. Table 4. Genes that are upregulated or downregulated in *Yap1* KD PS cells** (cutoff: fold change=1.3)

**Supp. Table 5**. **The top 6000 genes based on regulatory potential (RP) from Cistrome-GO analysis of YAP1 ChIP-seq data**.

**Supp. Table 6-7. Genes that are activated or repressed by YAP1 identified by integration of**
