## Supplementary figures and images for "P4HA2-induced prolyl hydroxylation suppresses YAP1-mediated prostate cancer cell migration, invasion, and metastasis"

### Suppl. Figure 1

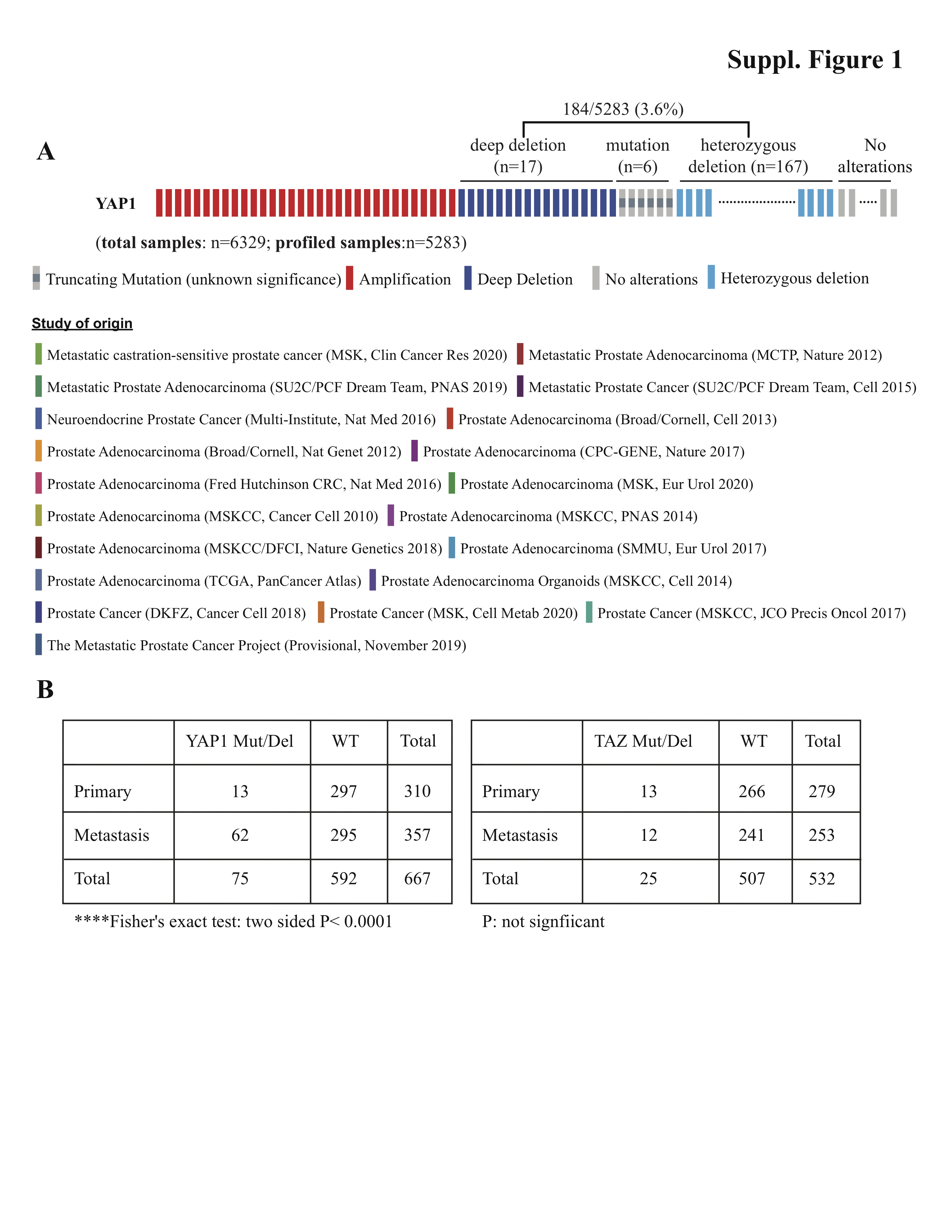

### Suppl. Figure 2

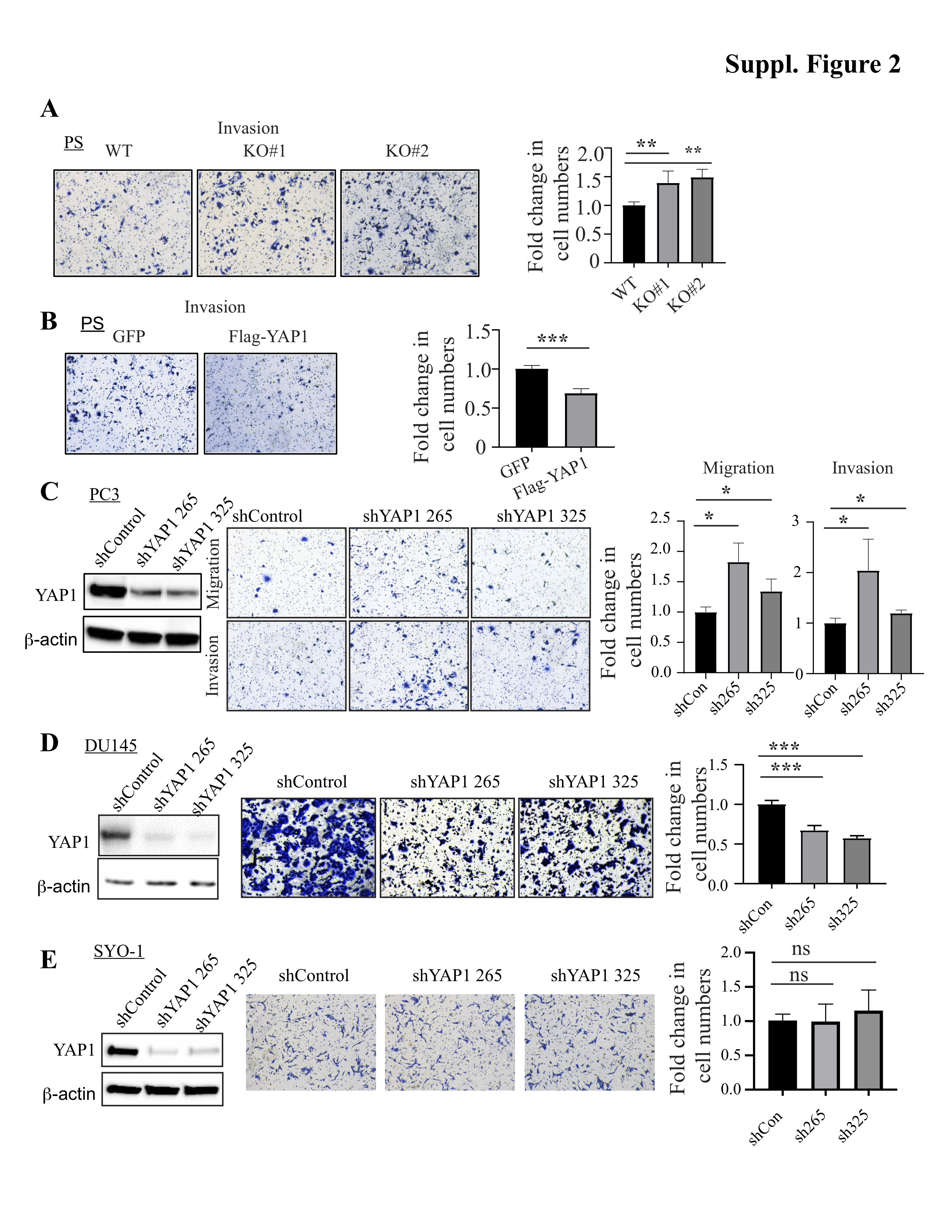

### Suppl. Figure 3

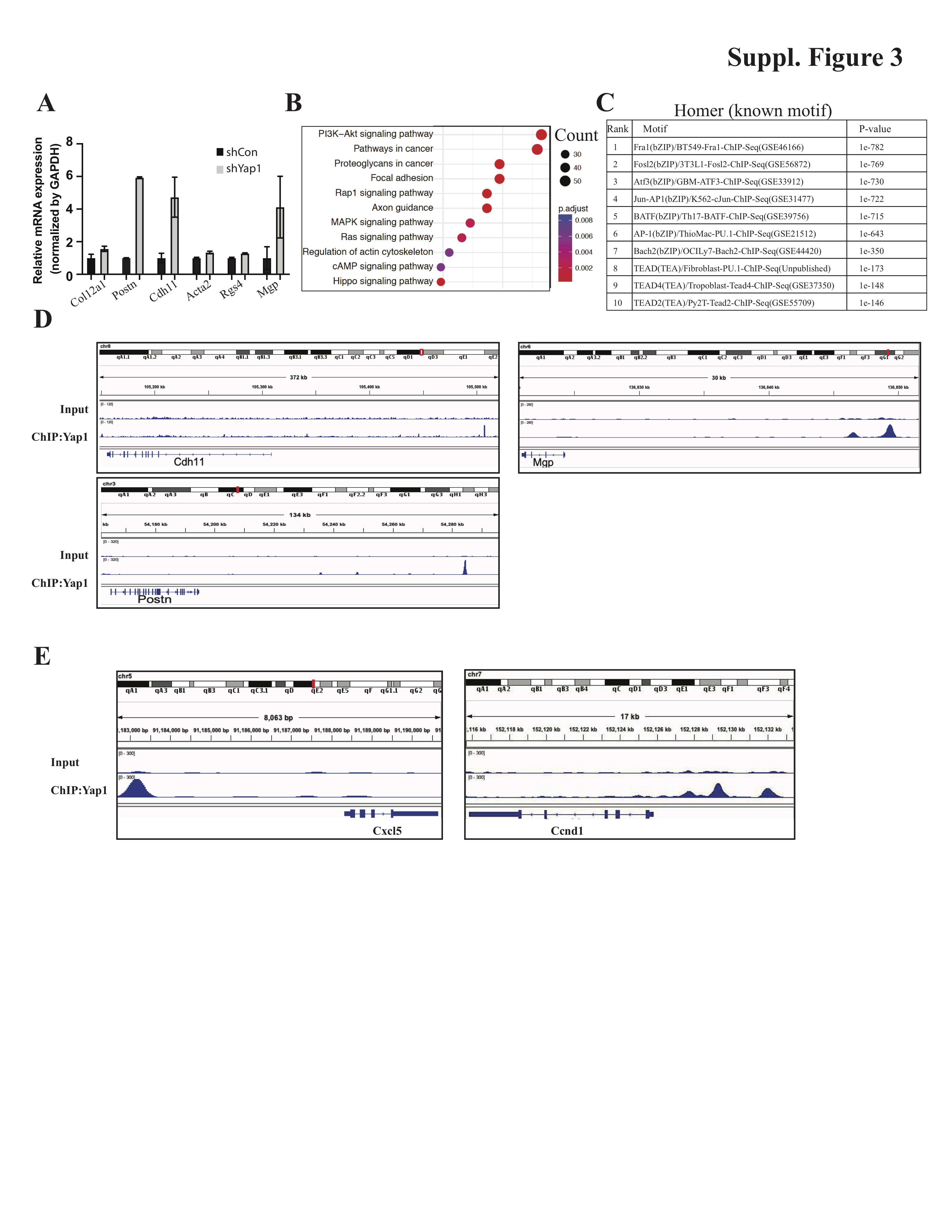

### Suppl. Figure 4

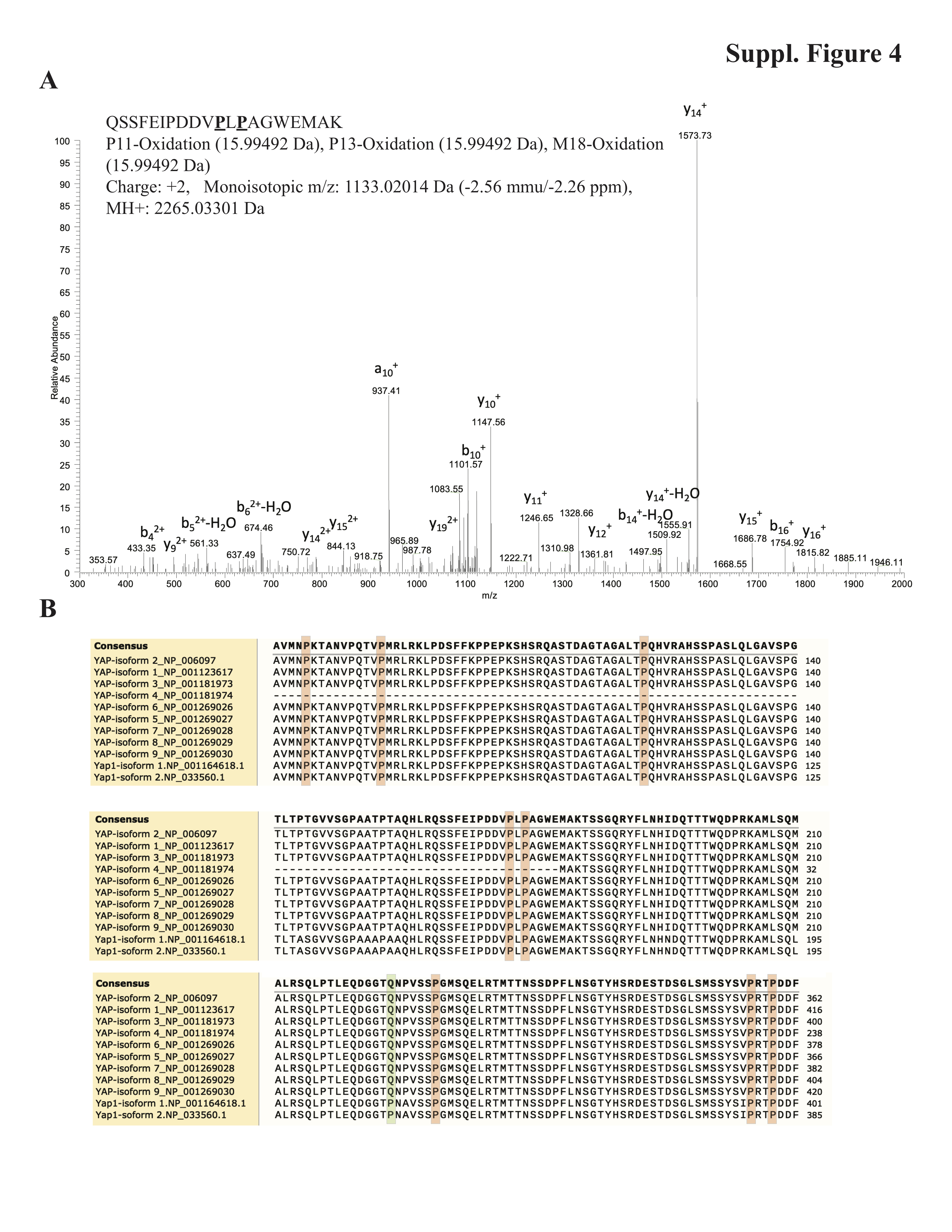

### Suppl. Figure 5

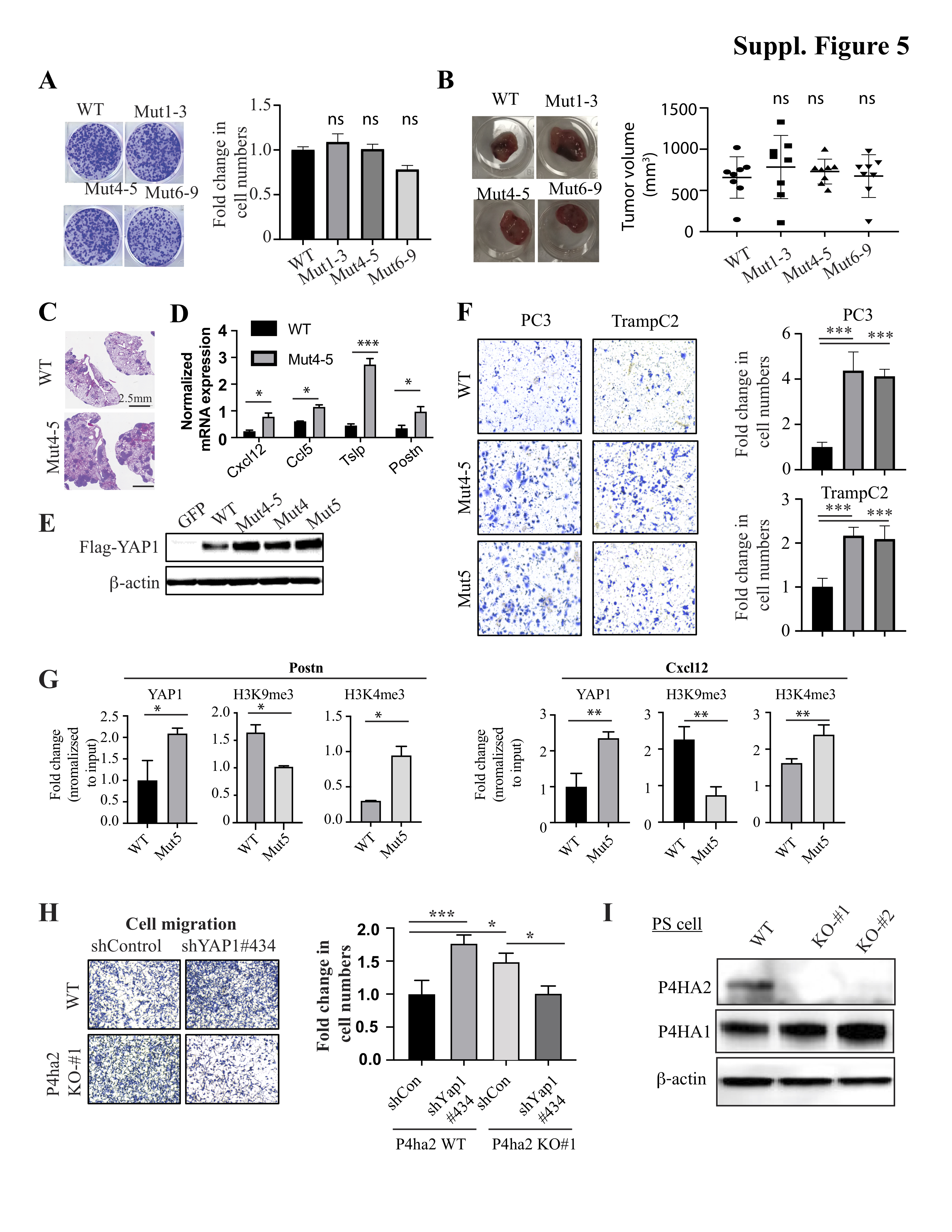
